# Investigation of hemodynamic bulk flow patterns caused by aortic stenosis using a combined 4D Flow MRI-CFD framework

**DOI:** 10.1101/2024.09.09.611958

**Authors:** Tianai Wang, Christine Quast, Florian Bönner, Malte Kelm, Tobias Zeus, Teresa Lemainque, Ulrich Steinseifer, Michael Neidlin

## Abstract

Aortic stenosis (AS) leads to alterations of supra-valvular flow patterns. These patterns might lead to, inter alia, increased damage of red blood cell (RBC) membranes. We investigated these patient specific patterns of a severe AS patient and their reversal in healthy flow through a 4D Flow MRI-based CFD methodology. Computational models of subject-specific aortic geometries were created using in-vivo medical imaging data. Temporally and spatially resolved boundary conditions derived from 4D Flow MRI were implemented for an AS patient and a healthy subject. After validation of the in-silico results with in-vivo data, a healthy inflow profile was set for the AS patient in the CFD model. Pathological versus healthy flow fields were compared regarding their blood flow characteristics, i.e. shear stresses on RBCs and helicity. The accuracy of the 4D Flow MRI-based CFD model was proven with excellent agreement between in-vivo and in-silico velocity fields and R² = 0.9. A pathological high shear stress region in the bulk flow was present during late systole with an increase of 125 % compared to both healthy flow. The physiological bihelical structure with predominantly right-handed helices vanished for the pathological state. Instead, a left-handed helix appeared, accompanied by an overall increase in turbulent kinetic energy in areas of accumulated left-handed helicity.

The validated 4D Flow MRI-based CFD model identified marked differences between AS and healthy flow. It suggests that altered turbulent and helical structures in the bulk flow are the cause for increased, potentially damaging forces acting upon RBCs in AS.

**Author Summary:** Aortic stenosis (AS) is a condition that alters the flow of blood through the aorta, potentially increasing damage to red blood cells (RBCs). It is therefore crucial to understand the precise nature of these changes, especially in the bulk region where the RBCs are mainly circulating. To investigate this, we developed a computational model based on patient-specific 4D Flow MRI data to simulate blood flow in a patient with severe AS and a healthy individual. Our high-fidelity models show high agreement with real-world data. We then imposed synthetically generated healthy flow conditions in the AS patient to eliminate age- and geometry-related uncertainties. Our results revealed significantly higher shear stresses and changes in helical flow patterns in the AS patient compared to healthy flow, suggesting that these alterations could increase the risk of RBC membrane damage. These findings underscore the importance of deepening our understanding about the interplay between bulk flow alterations and their impact on RBCs, as previous studies were mostly focused on near wall regions. This methodology could also be extended to other cardiovascular conditions, aiding in the development of patient-specific treatment strategies.

## Introduction

The development of cardiovascular diseases is linked to altered hemodynamic environments of the volumetric blood flow and therefore, gaining a deeper insight into both pathological and physiological flow conditions is of undeniable importance [1, 2]. A pathology of high prevalence and clinical relevance is aortic valve stenosis (AS) and the American Heart Association (AHA) predicted an increase of more than 100 % in patients suffering from this disease by 2050 [3]. A better understanding of AS pathophysiology can help to develop safer and more reliable therapies to tackle these upcoming challenges. On this front, aortic flow patterns during AS play a decisive role in the pathology [4]. Valvular pathologies lead to disturbed blood flow in the aorta which in turn has the potential to damage red blood cell (RBC) membranes. In order to capture these in-vivo flow features non-invasively, 4D Flow Magnetic Resonance Imaging (4D Flow MRI) is commonly used. 4D Flow MRI alone, however, does not provide sufficient temporal and spatial resolution due to its space- and phase-averaging nature to reliably derive all velocity scales and gradient-based hemodynamic parameters, such as shear stress fields on circulating particles [5], which are essential quantities to investigate the cause of blood damage [2, 6].

In order to overcome this limitation, patient-specific Computational Fluid Dynamics (CFD) has been gaining popularity in the field of medical engineering research. Here, the advantages of high spatial and temporal resolution of CFD simulations are combined with subject-specific information about geometry and boundary conditions. Different approaches have been developed over time to increase patient-specificity and therefore clinical credibility of these models, e.g. the use of realistic geometries segmented from medical images. More recent research is focusing on the definition of accurate boundary conditions at the inlets and outlets of vessel [2]. These boundary conditions can be defined in varying degrees of complexity and accuracy. Some studies used generic curves and uniform or parabolic inlet waveforms [7–10], or, similarly, simplified conditions at the outlets, such as constant or waveform pressure [7–9, 11]. One-dimensional modeling strategies at the inlet, however, neglect the spatial in-plane distribution of velocity which is highly complex and particularly characteristic and therefore essential for non-physiological flow states, such as AS [12]. Pirola et al. [13] emphasize the importance of including secondary flows in the prescription of inlet profiles in order to achieve realistic velocity, stress and helicity fields, in particular in patients with aortic valve pathologies. Highest subject-specificity and agreement with in-vivo flow can be achieved by using flow measurement techniques, such as Doppler echocardiography, time-resolved 2D Flow MRI or 4D Flow MRI. While Doppler echocardiography can only provide one-directional flow information in two dimensions and the current gold standard for volumetric flow analysis of 2D Flow MRI is limited by a two-dimensional imaging plane, 4D Flow MRI is known to be the only existing method for non-invasive, time-resolved, and three-dimensional measurement of blood velocity in-vivo [2, 7].

When it comes to comparing patient-specific flow patterns, the definition of a control group poses challenges. Using age-matched individuals introduces anatomical differences as an unavoidable confounding factor. Furthermore, it remains challenging to recruit age-matched scans of healthy volunteers without any other diseases or pathologies [14]. According to the AHA, about 77-91 % of the American population aged > 60 years suffer from a cardiovascular disease [15]. A potential approach to overcome this limitation could be the generation of a patient-specific synthetic healthy model as in-silico physiological reference for a patient, using pre-acquired knowledge about physiological flow profiles and patterns. According to Youssefi et al. [12], patient-specific profiles can be sufficiently substituted by idealized profiles when investigating healthy flows.

In summary, investigations of pathological aortic bulk flow structures during AS, and their relationship to potential RBC damage are lacking. In the past, the focus has mostly lied on the near-wall hemodynamics, e.g. wall shear stresses, residence times and pressure distributions, and only little knowledge is available on volumetric bulk properties within the vessel lumen. However, bridging this gap is indispensable to continuously deepen the clinical knowledge and understanding of in-vivo aortic hemodynamics and their interplay with cardiovascular diseases.

Therefore, the goal of the present work is to develop a 4D Flow MRI-based CFD methodology to generate patient-specific models of healthy and pathological aortic flow and to show its ability to accurately characterize blood bulk flow structures and alterations caused by AS in the aortic arch. Applying this methodology to large patient cohorts and by comparing healthy and pathological flow structures, hemodynamic differences between these flow fields will aid the identification of those flow features with potentially damaging effect on circulating erythrocytes during AS.

## Materials and Methods

The complex aortic flow fields of a healthy adult and an adult suffering from severe AS were determined using a methodology of CFD analysis based on 4D Flow MRI data and are described in detail in the following. The two datasets served as input data for the generation of a synthetic healthy model of the AS patient. The derived hemodynamics were compared using qualitative and quantitative methods to identify the largest differences in the aortic flow structures between the physiological and pathological conditions measured in this study.

### Study design

4D Flow MRI and multisliced computed tomography (CT) measurements of a 78 year-old, female patient diagnosed with severe AS (AS-78), originally acquired at the University Hospital Düsseldorf (UKD) were anonymized and made available for retrospective analysis. In addition, a 4D Flow MRI scan of a healthy (H-25) volunteer (female, age = 25 years) was acquired at the University Hospital Aachen (UKA) as physiological reference. Approvals were granted for all acquisitions by the respective Institutional Review Boards. 4D Flow data were acquired on 1.5 T MRI systems (Achieva at UKD and Ingenia Ambition at UKA, Philips Healthcare, The Netherlands) using the torso and posterior coil. Relevant scan parameters are listed in Supplement I (S-I). Single source CT data were acquired using a SOMATOM Definition AS+ (Siemens Healthcare, Forchheim, Germany), with a temporal resolution of 150 ms and collimation of 128 x 0.6 mm for the AS-78 patient.

### In-vivo data processing

The handling of in-vivo data followed general guidelines and recommendations presented in the cardiovascular 4D Flow MRI consensus statement by Dyverfeldt, Bissell et al. [2]. The phase contrast DICOM images were processed using in-house developed MATLAB R2021b (MathWorks, Natick, MA) scripts. The data were first sorted according to timeframe and slice number.

Furthermore, the stored greyscale values were transformed to their flow values in cm/s by multiplication with the velocity encoding value (Venc). The magnitude images were used to segment the aortic geometry (aortic arch and supraaortic vessels) during systolic peak ejection with the open-source software ITK Snap (www.itksnap.org) [16]. For the AS case, the segmentation was done manually on a layer by layer basis by tracking the outer contour of the aortic arch. The magnitude images of the healthy volunteer provided sufficient signal-to-noise ratio and grey value contrast to enable semi-automated segmentation of all aortic parts, including the supra-aortic vessels, by interpolation between adjacent slices. Finally, the magnitude-based binary segmentations were superposed with the volumetric velocity data for each timeframe from the 4D Flow MRI scans using an in-house MATLAB script. This yielded a segmented velocity volume for the patient-specific lumen of the thoracic aorta. Resulting flow fields were visualized in ParaView 5.10.1 (KitWare, Clifton Park, NY).

### Computational model

For subsequent generation of the numerical models, the commercially available CFD software package ANSYS CFX 2021 R2 was used, including the solid modeling Computer-Aided Design (CAD) software ANSYS SpaceClaim 2021 R2 and the meshing tool ANSYS ICEM CFD 2021 R2, all developed by ANSYS, Inc. (Canonsburg, PA).

The vascular geometry for the AS-78 case was semi-automatically segmented from CT images of the same patient. For the healthy case, the segmentation obtained from the 4D Flow MRI scans was used. Subsequently, the segmentation surfaces were manually smoothed using ANSYS SpaceClaim and all inlet and outlet surfaces were cut perpendicularly. Finally, the patient-specific geometries were meshed with suitable grid settings ensuring mesh independency of the numerical results. Meshing was performed with ANSYS ICEM CFD using unstructured tetrahedral meshes and prism elements for the near wall regions. A mesh independence study (see Supplement II (S-II) for further information) yielded a number of 4.65 million elements and 12 prism layers with a refinement region in the ascending aorta. Subsequently, the meshes were imported to ANSYS CFX. Here, the numerical model was built using patient-specific boundary conditions derived from the in-vivo measurements.

The Carreau-Yasuda model was implemented for modeling of the Non-Newtonian blood behavior in terms of a shear rate γ dependent viscosity η function, utilizing parameter values defined by Abraham et al. [17] and a constant density *ρ* of 1060 kg/m^3^.

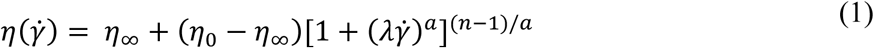

Here, η_0_is the viscosity when the shear rate tends to zero, η_∞_ is the viscosity for high shear rates, λ is a time constant, *a* and *n* are dimensionless parameters. Turbulent flow has been observed in AS in numerous studies, e.g. in a turbulence analysis by Manchester et al. [18]. According to Benim et al. [19], the SST model can efficiently adapt to transitional effects and is widely used in industrial and academic context. Therefore, the Reynolds-averaged SST model was chosen to describe the expected formation of turbulent eddies.

The inlet of the numerical model was located shortly above the aortic valve and the 4-dimensional velocity profile serving as input data was extracted from the corresponding plane in the 4D Flow MRI velocity volume. The velocity values were interpolated linearly in both space and time to adjust to the high-resolution CFD grid and time using an in-house MATLAB script. A preliminary analysis of cycle-dependency showed, that the results converged for the third heart cycle. Thus, three cardiac cycles were simulated and the third one was post-processed, with a numerical time step size of 1/40 of the corresponding MRI timeframe.

The supra-aortic vessels and the descending aorta were treated as openings defined by loss coefficients, which control the resulting mass flow in the respective branch and serve as a substitute of the downstream vasculature and its impact on the flow structure. These loss coefficients can only be determined iteratively. Thus, for each numerical time step, an inner feedback loop was implemented with 100 iterations, which compared the percentage mass flow distribution through one opening for each iteration step with a prescribed percentage flow. According to the deviation, the loss coefficient was adapted for the next iteration until the two values were synchronized. This process was repeated for each numerical time step. The idealized percentage flow distributions were calculated from values provided in Benim et al. [19].

Finally, the vessel wall was modeled as a rigid, no-slip wall. Convergence control was set for a root mean square residual level of 10e-5 for the mass and momentum conservation equations.

### Synthetic healthy model

Due to the differences in age and geometry of the H-25 and the AS-78 subjects, a synthetic healthy model of the AS patient was developed and checked for credibility by comparing the resulting flow patterns with the healthy model. For the development of the synthetic healthy model of the AS patient (H-78), a parabolic profile was assumed, which, according to Youseffi et al. [12], is sufficiently accurate as inlet conditions to be a simplified substitute for medical data-derived profiles in case of physiological flow. The parabolic profile was parameterized and adapted to match the 4D Flow MRI-derived volume flux of the AS patient at each MRI acquisition timestep. The resulting geometry-specific, parabolic velocity profiles were interpolated between these time points by fitting a Fourier series. According to Morbiducci et al. [20], in-plane velocity components are non-negligible for bulk flow structure investigations, adding up to approx. 32 % of axial velocity during systole and 111 % during diastole. Therefore, secondary flows were prescribed as percentage amounts of the current axial flow magnitude. The percentage values were derived from the healthy case H-25 for each MRI timestep and used to determine the Fourier series coefficients to fit these datapoints as well. This resulting patient-specific, three-dimensional velocity waveform was prescribed as inlet boundary condition to mimic synthetic healthy flow of the AS patient. The detailed calculations and Fourier series fittings can be found in Supplement III (S-III). In total, three CFD models were created and the setup is summarized in Table 1.

**Table 1.** Overview of the three CFD models generated within the scope of this study. H: healthy, AS: aortic stenosis. The age is provided in years.

| Model ID | State | Age and gender | Inlet condition | Geometry | Purpose |
| --- | --- | --- | --- | --- | --- |
| H-25 | H | 25, female | 4D Flow MRI | H | Credibility check for H-78 |
| AS-78 | AS | 78, female | 4D Flow MRI | AS | Numerical model used to elucidate flow alterations in AS |
| H-78 | H | 78, female | 4D synthetic parabolic | AS | Synthetic healthy model for specific AS patient, to eliminate age-, geometry and other individual-related influence factors, and to overcome the lack of data availability |

### Numerical evaluation

Finally, several parameters of interest were extracted from the CFD models for the validation of the model as well as for the subsequent investigation of the aortic flow behavior at characteristic timeframes. The following cross-sectional and sagittal planes served as regions of interest (ROI) for investigation: ROI A, ROI B, ROI C and SAG, as shown in Fig. 1.

**Fig. 1.**
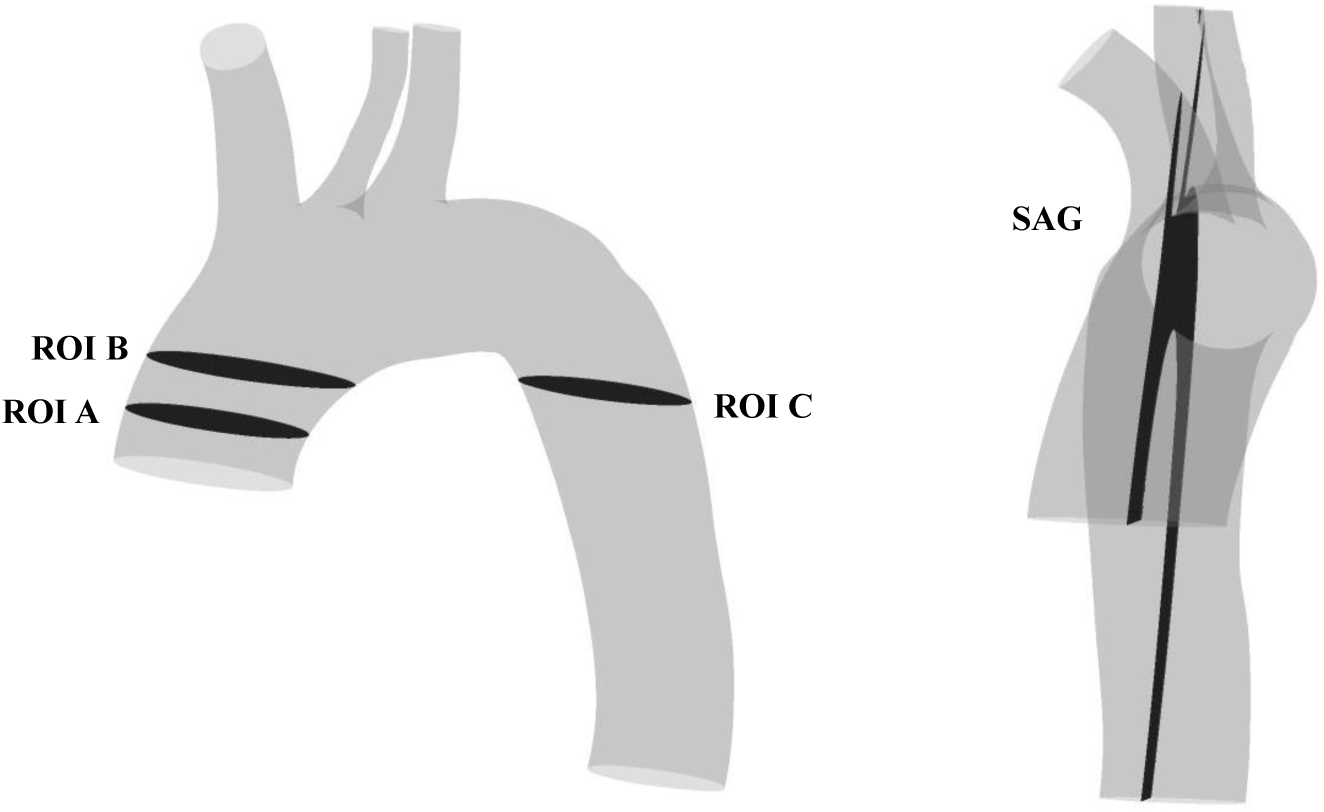
Location of ROIs in the aortic model which are used for validation and post-processing of the numerical results

The temporal and spatial velocity fields were subjects of investigation for validating the generated CFD model. Therefore, the area-averaged velocity values on ROI A-C were compared between the CFD and MRI data for all MRI timeframes by means of linear regression analysis. Further, spatial velocity distribution within these ROIs was analyzed qualitatively for the systolic timeframe.

In order to locate the region of highest forces in the bulk flow acting upon erythrocytes potentially causing damage for the AS case, acting stresses and turbulence parameters were processed in both their temporal and spatial properties. The following quantities were of interest: Total shear stresses (TSS), together with Reynolds shear stresses (RSS) and viscous shear stresses (VSS). Turbulent kinetic energy (TKE), helicity and local normalized helicity (LNH). Their individual derivation and implementation is explained in the Supplementary Material (S-IV).

## Results

### Validation of the in-silico model

For a quantitative evaluation of the temporal evolution of velocity predicted by the CFD simulation, the area-averaged velocity at ROI A, B and C are plotted over the cardiac cycle for MRI and CFD for H-25 and AS-78, respectively. The temporal evolution for ROI B and C are shown in Fig. 2 a, b, d and e, the corresponding evolution for ROI A can be found in the Supplementary Material (S-V). The noise of each acquisition was calculated by determining the standard deviation of the measured velocity in the surrounding non-vessel regions during diastole, where no signal should exist. The resulting noise values of approx. 57 mm/s for AS-78 and 46 mm/s for H-25 correspond to approx. 5% of the Venc value, which Papathanasopoulou et al [21] determined to be an approximation of the noise in MRI measurement. The noise is represented by the shaded areas in Fig. 2. Hence, the numerical results show very good agreement concerning the shape and magnitude of the area-averaged velocity. Additionally, a linear regression analysis for ROI A-C and SAG deliver an R^2^-value of 0.9 (p < 0.001) for both the physiological and the pathological flow, indicating excellent agreement and statistically significant correlation between 4D Flow MRI-based CFD results and the measured in-vivo 4D Flow MRI velocities. The corresponding regression plots are presented in Fig. 2 c and f.

**Fig. 2.**
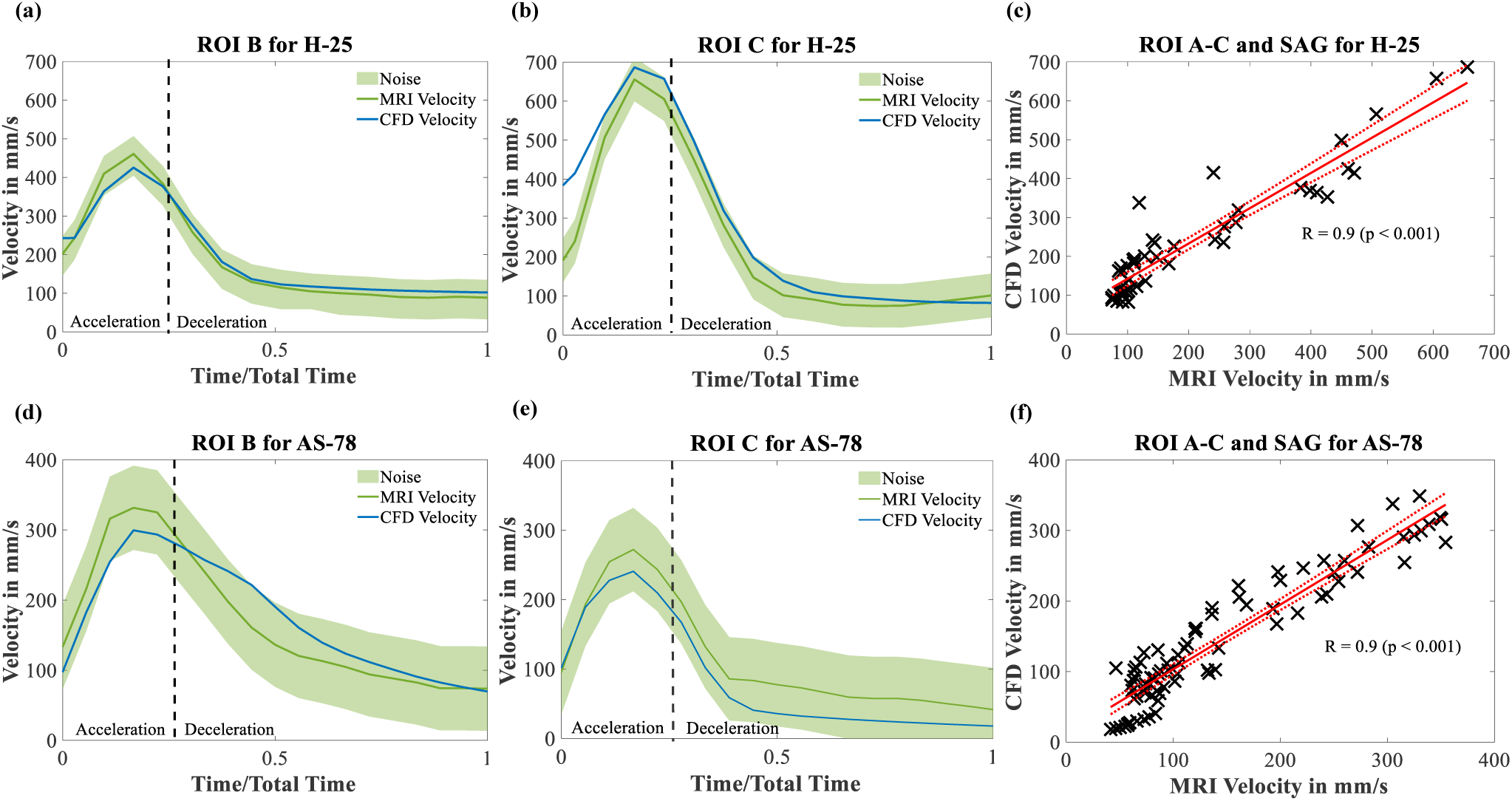
Validation results of the 4D Flow MRI-based CFD model, each plot compares the 4D Flow MRI measured values with the corresponding numerically determined velocities. (a-b) Area-averaged temporal evolution of velocity for H-25 case on ROI B and C, respectively. (c) Linear regression plot for H-25 velocities on ROI A-C and SAG for area-averaged velocities throughout the cardiac cycle. (d-e) Area-averaged temporal evolution of velocity for AS-78 case on ROI B and C, respectively. (f) Linear regression plot for AS-78 velocities on ROI A-C and SAG for area-averaged velocities throughout the cardiac cycle. Abbreviations: H-25 – Healthy, 25 year old volunteer, AS-78 – Aortic Stenosis, 78-year old patient, ROI – region of interest

Further, the numerically calculated and measured velocity fields also show high agreement spatially. Examples of spatial flow patterns and depictions of momentary regions of higher and lower velocities on ROI A-C are appended in the Supplement Material VI (S-VI) for the pathological flow field during early systole. Here, the difference between the high resolution CFD grid and the corresponding low-resolution MRI grid also becomes obvious.

### Physiological and pathological in-silico velocity fields

With excellent agreement between the in-vivo and in-silico results, the in-silico velocity fields are investigated in more detail for the three numerical models. First, a credibility comparison between H-25 and H-78 shows that both models reproduce an overall homogeneous, parabolic shaped and rotationally symmetric velocity distribution in the ascending aorta, as shown in Fig. 3. In the descending aorta, however, large differences can be observed with high velocities for H-25, while the velocity magnitudes remain relatively constant along the aortic arch for H-78.

**Fig. 3.**
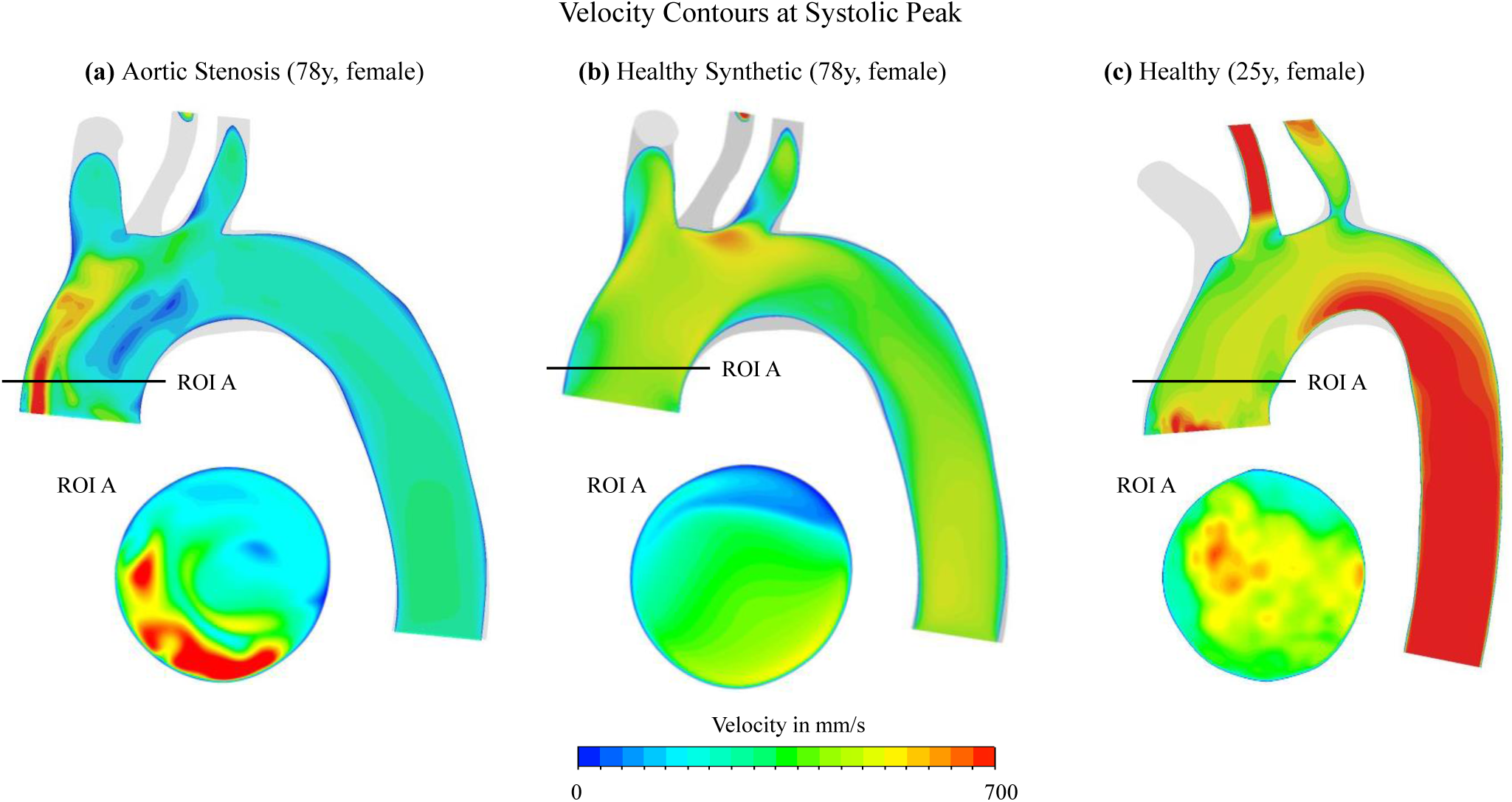
Velocity contours on sagittal and cross sectional plane ROI A at systolic peak for a) aortic stenosis case AS-78, b) synthetic healthy case H-78 and c) healthy case H-25

In contrary, the numerically derived systolic aortic stenotic flow displays an eccentric high velocity jet flow along the outer wall of the ascending aorta. When comparing the maximum and averaged velocities of cross-sectional planes in the ascending aorta over a cardiac cycle, the area-averaged systolic value is higher in the healthy cases, whereas the peak velocity is significantly higher for the aortic stenosis case, due to the stenotic jet. In the descending aorta, both average and maximum velocity are higher in physiological flow. Thus, the highest velocities are located in the ascending aorta for the AS patient and further downstream for both H-25 and H-78. The volume-averaged (maximum) Reynolds number (Re) at systolic peak in the ascending aorta is Re = 2350 (5126.5) for AS-78, Re = 2027 (7887.19) for H-78 and Re = 2447 (6950.8) for H-25.

### Shear stresses acting onto erythrocytes

First, the total shear stress (TSS), viscous shear stress (VSS) and Reynolds shear stress (RSS) are compared between the healthy and the synthetic healthy flow. The synthetically generated shear stress field on ROI A displays similar temporal and spatial distribution as the reference field in H-25. In both cases, RSS remains negligible and the TSS and VSS peak coincides with the moment of systolic velocity peak, as shown in Fig. 4. The peak TSS contour on ROI A is rather homogeneous in both cases with the highest values occurring at the vessel wall. Further, Fig. 4 depicts results of the pathological flow on ROI A. The area-averaged, temporal evolution shows, that TSS for the AS-78 case is significantly higher than for the H-25 and H-78 case, with a second peak occurring during deceleration and being the global stress peak. At this point, it becomes 125 % higher than the TSS value in the healthy case. The first shear stress peak also shifts into the deceleration phase whereas it occurred at the systolic peak for both healthy cases. Further, RSS as an indicator for turbulent flow is now clearly contributing to the overall acting stresses for AS. The AS-78 TSS contour in Fig. 4 shows the eccentric region of high TSS in the pathological bulk flow at its peak, whereas the physiological stress fields do not show any obvious characteristics at its respective peak.

**Fig. 4.**
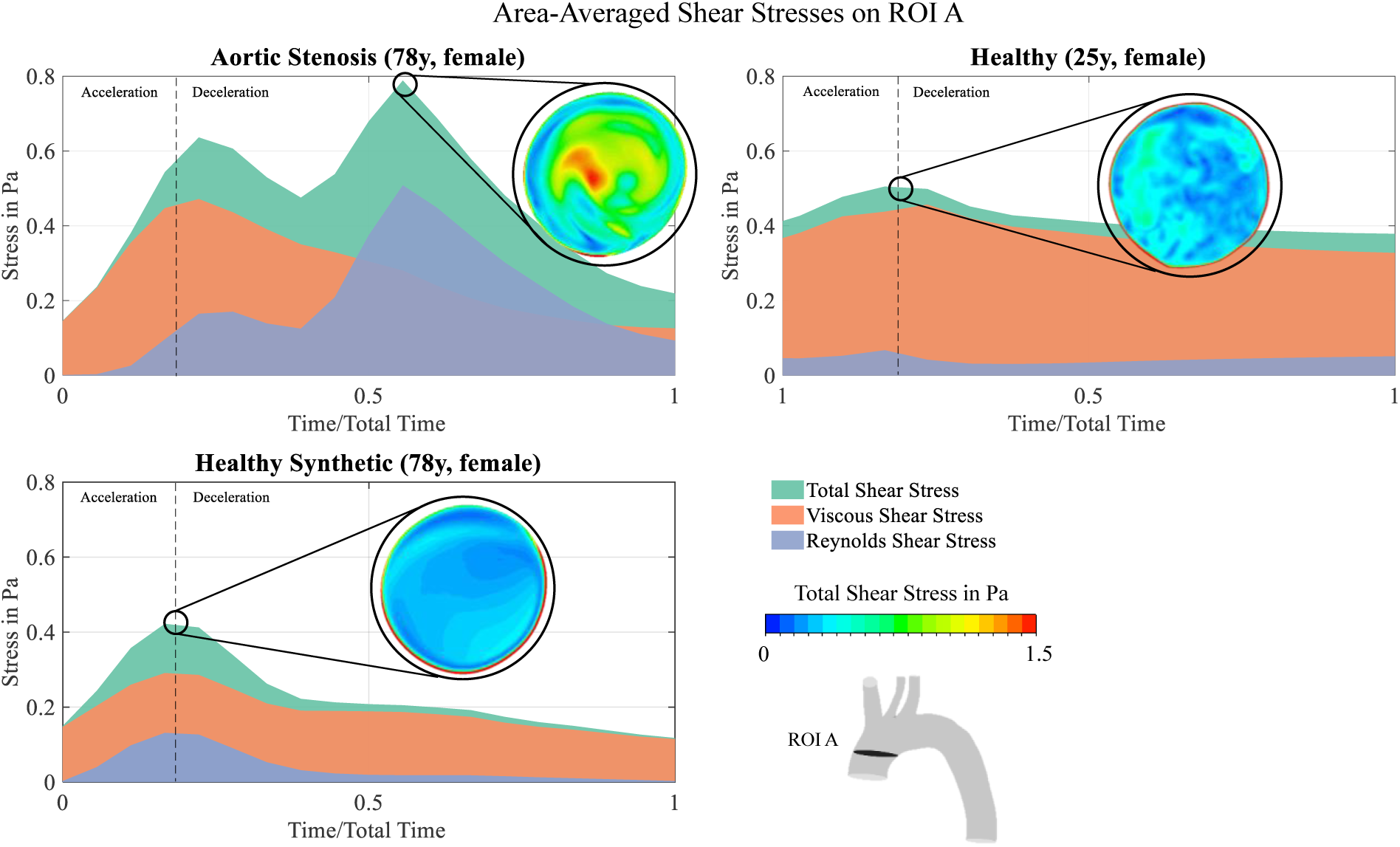
Temporal area-averaged shear stress evolutions and contours at their respective peak time frames on ROI A. Upper left: Shear stresses for the aortic stenosis case AS-78. Upper right: Shear stresses for the healthy case H-25. Bottom left: Shear stresses for the synthetic healthy case H-78.

With the significant rise in bulk RSS for AS-78, bulk flow behavior is investigated in detail. First, turbulent kinetic energy (TKE) as a quantity of turbulence is compared qualitatively on the sagittal plane SAG. Both H-25 and H-78 display negligible elevations of TKE values in the entire domain, with a small region of higher TKE in the central aortic arch, see Fig. 5. The analysis shows strong alteration for AS flow with swirling structures in the ascending aorta.

**Fig. 5.**
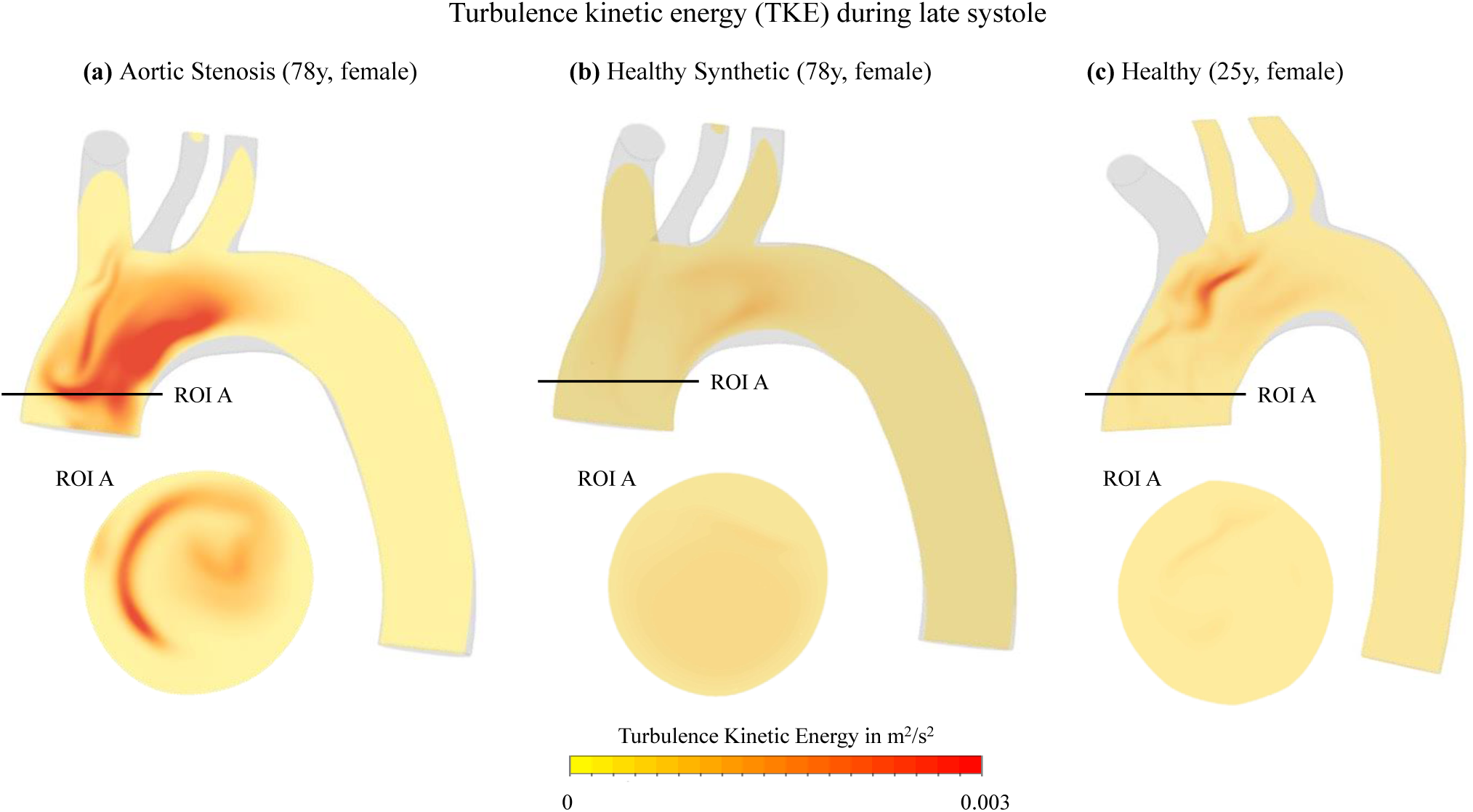
Turbulent kinetic energy distribution on sagittal plane SAG during deceleration for (a) aortic stenosis case AS-78, (b) synthetic healthy case H-78 and (c) healthy case H-25

Second, helicity and its locally normalized value, namely Local Normalized Helicity (LNH), were analyzed in the ascending aorta over the heart cycle and during late systole, respectively, as shown in Fig. 6. For both healthy models, the average helicity values remain positive over the entire cardiac cycle with their peak occurring shortly after the systolic velocity peak in ROI A and B. The area-averaged values are lower for H-78 and the cross-sectional contour of the synthetic healthy case displays less chaotic patterns and a smooth transition between a smaller dominantly left-handed to a larger dominantly right handed region, whereas the H-25 case shows more mixed patterns. Overall, if putting a convex hull around the predominantly right- and left-handed regions, both cases lead to similar distributions of positive and negative helicity areas. In AS, however, the mean helicity turns from right-handed to left-handed characteristics during late systole, and the magnitude reaches its peak later than in the healthy case. Fig. 6 c and d present the LNH contours on ROI A at their peak during deceleration. The healthy flow displays two counterrotating helices (b, c), while one nearly central, left-handed vortex can be identified for the AS flow (a). When comparing the LNH contour for AS-78 with the corresponding TKE contour in Fig. 5, the region of highest TKE coincides with the region of left-handed helicity (LNH < 0).

**Fig. 6.**
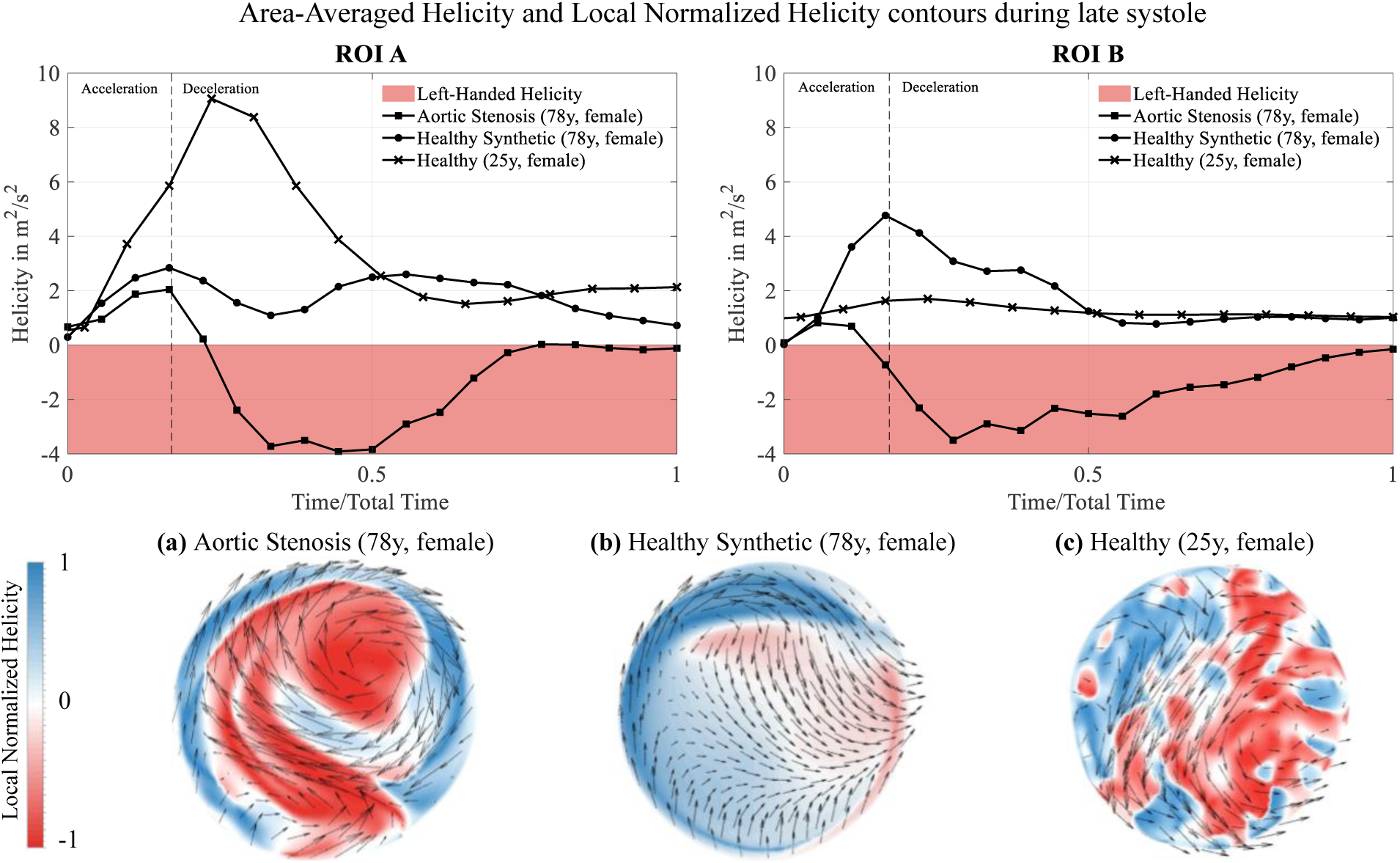
Temporal and spatial helicity distribution. Upper row: Area-averaged helicity evolution on ROI A and B for AS-78, H-78 and H-25. (a) LNH contour and tangentially projected streamline vectors on ROI A during deceleration peak for AS-78. (b) LNH contour and tangentially projected streamline vectors on ROI A during deceleration peak for H-78. (c) LNH contour and tangentially projected streamline vectors on ROI A during deceleration peak for H-25.

## Discussion

The present study aimed to identify the alterations in aortic blood flow patterns caused by aortic valve stenosis which may lead to erythrocyte membrane damage. This was achieved by developing a subject-specific and realistic numerical flow model using in-vivo 4D Flow MRI. After validating the CFD model results with in-vivo data of two subjects, differences between AS and healthy aortic bulk flow were analyzed. In order to omit confounding factors such as anatomy and age, a synthetic healthy profile was set for the AS patient geometry. Especially for healthy flows, such approach is justifiable, as outlined by Youssefi et al. [12]. In order to create more realistic flow conditions, we enhanced the synthetic flow profile by the same secondary flow components as seen in the healthy subject of 25 years (H-25). This aspect is crucial when one aims to realistically model vorticity and helicity of aortic flow [20, 22].

Our comparison of the synthetic healthy (H-78) and realistic healthy (H-25) yielded overall good agreement in ascending aorta flow features such as homogeneous velocity distributions, very low turbulent kinetic energies and right-handed helices (see Fig. 3-6). With the latter flow feature being underestimated by the synthetic flow case. Differences in the descending aorta with H-25 flow velocities being higher than H-78 flow velocities can be attributed to age-dependent changes. Interestingly, the in-vivo study of Garcia et al. [23] has identified similarly increased flow velocities in the descending aorta when comparing female subjects of 21-39 years with subjects of > 60 years. Overall, the synthetic healthy flow model H-78 can be considered sufficiently credible to serve as patient-specific, healthy reference for the AS patient in our study and was used for the subsequent comparison with the AS flow. Thus, in the remaining analysis pathological (AS-78) and healthy (H-78) were compared.

### Flow alterations during aortic stenosis and erythrocyte damage

Differences in the cross-sectional velocity distribution were caused by the presence of a high-velocity jet due to the reduced orifice of the aortic valve in AS. In order to pinpoint the impact of aortic flow alterations on shear stress on circulating cells (e.g. erythrocytes), shear stress distributions, turbulence kinetic energy and helicity were investigated in detail. The TSS evolution displays a key novel finding: it develops a second peak for the pathological flow during early diastole, which shows an increase by 125 % compared to the corresponding maximum physiological values. Therefore, the increased shear stresses acting in the bulk flow, which are indicative of the pathological state during AS, are likely to impact and apply damaging forces on circulating RBCs. Nonetheless, only the spatial characteristic of forces acting onto the erythrocytes is investigated in this present work. Flow-induced damage of erythrocytes, however, is an integrated effect of both spatial and temporal factors and Lagrangian-based analyses are required, since the instantaneous streamlines and particle pathlines do not align in complex aortic flow.

Next, an additional investigation of the bulk flow structures during the TSS peak was carried out for deeper understanding. Here, an increase in turbulence characteristics in the bulk flow of the ascending aorta is present in AS, whereas TKE was negligible for the healthy case. Further, significant differences in helicity were identified for the ascending aortic bulk hemodynamics, which is a known parameter to evaluate blood flow alterations, particularly caused by valvular diseases [22, 24]. However, only little is known about the helical properties during AS [25]. According to Kilner et al. [26] and Markl et al. [27], two consistent features present in physiological flow of the human aorta are: helicity and retrograde flow. The anatomical curvature of the aortic arch leads to right-handed, well-aligned helical outflow during systolic peak. During deceleration, the ascending aorta is still predominantly filled with right-handed helices with a small portion of retrograde flow developing over a short distance during end-systole, which Morbiducci et al. [22] describe as a bihelical pattern with two counterrotating, Dean-like vortices, which is also characteristic for fully developed flow in a bended pipe. This can also be detected in the healthy flow structure of H-25 in this study, and to some extent in H-78, given its nature to underestimate helicity as described earlier. The numerical results of the pathological flow in this study, however, show a decrease in right-handed helicity and an increase of the region of left-handed vortices. These areas of accumulated left-handed helices coincide with the areas of high TKE (see Fig. 5 and Fig. 6), thus pointing towards a correlation between left-handed helices and high presence of RBC-damaging turbulence potentially impacting erythrocyte function.

Overall, the results allow for the formulation of two main findings that – to the authors’ best knowledge – have not been identified in existing literature so far:

1. The presence of a second TSS peak in the bulk flow during early diastole was observed, which correlates with the onset of strong turbulent structures in AS flow.
2. AS further causes a decrease in physiological right-handed helical structures. This also leads to the destruction of the bihelical structure, which is found to be characteristic for physiological aortic flow. Instead, a main left-handed helix is present, coinciding with the peak region of TKE.

These two hemodynamic alterations are hypothesized to cause damage on circulating cells and cellular dysfunction. However, our work only included a size of N = 2. Therefore, these findings need to be further validated with a larger cohort of patients with the developed framework.

### 4D Flow MRI-based CFD methodology

In order to validate the developed framework and the resulting findings, the contours of the numerical velocities profiles were analyzed on a set of planar, cross-sectional ROIs along the aortic centerline. Their temporal and spatial agreement with the corresponding velocity profiles obtained in the MRI process were compared qualitatively and quantitatively. A good qualitative agreement of patterns and contours is found on the ROIs. The linear regression analysis used to assess the overall correlation between the measured MRI velocity and the predicted CFD results yields excellent results with R^2^ = 0.9 for both the physiological and the pathological case.

To the best of the authors’ knowledge, this is the first study to compare patient-specific bulk flow patterns in healthy flow and aortic stenosis by combining spatially and temporally resolved 4D Flow MRI measurements and CFD modeling in order to identify the pathological impact on erythrocytes due to flow alterations during AS. It implements realistic, four-dimensional in-vivo boundary conditions and the numerical modeling of non-Newtonian and turbulent blood behavior. However, the 4D Flow MRI acquisition and post-processing suffered from limitations characteristic for this technique, e.g. the averaging nature of the scans, systematic errors [2], or segmentation uncertainties. Further, simplifying the vessel wall with rigid properties can lead to overestimations of some quantities such as the wall shear stress. According to Torii et al. [28], however, this overestimation becomes negligible (< 5 %) when analyzing time-averaged values and was therefore accepted in this study to keep computational costs low.

Large differences between the given models of healthy flow and the altered hemodynamics present during aortic stenosis were identified: The occurrence of a second TSS peak in the bulk flow of the ascending aorta during early diastole and the loss of the physiological, predominantly right-handed bihelical structure. The left-handed helices coincided with areas of high TKE. These results were obtained by the development of a 4D Flow MRI-based CFD framework of high fidelity. These patient-specific models promise patient-individual treatment planning in the future, e.g. for aortic valve replacement, and improved long-term outcomes of patients undergoing such procedures.

## Notes

**Statements and Declarations: Funding:** This research project was supported by the Studienstiftung des Deutschen Volkes (PhD Fellowship, German Academic Scholarship Foundation) and the Deutsche Forschungsgemeinschaft (DFG, German Research Foundation) CRC/TRR259, Grant No. 397484323.

### Competing Interest Statement

The authors have declared no competing interest.

## References

1. Long Q, Xu XY, Ariff B, Thom SA, Hughes AD, Stanton AV. Reconstruction of blood flow patterns in a human carotid bifurcation: A combined CFD and MRI study. J. Magn. Reson. Imaging. 2000;11:299–311. doi:10.1002/(SICI)1522-2586(200003)11:3<299::AID-JMRI9>3.0.CO;2-M.

2. Dyverfeldt P, Bissell M, Barker AJ, Bolger AF, Carlhäll C-J, Ebbers T, et al. 4D flow cardiovascular magnetic resonance consensus statement. J Cardiovasc Magn Reson. 2015;17:72. doi:10.1186/s12968-015-0174-5.

3. Tsao CW, Aday AW, Almarzooq ZI, Alonso A, Beaton AZ, Bittencourt MS, et al. Heart Disease and Stroke Statistics-2022 Update: A Report From the American Heart Association. Circulation. 2022;145:e153–e639. doi:10.1161/CIR.0000000000001052.

4. Gülan U, Lüthi B, Holzner M, Liberzon A, Tsinober A, Kinzelbach W. An in vitro investigation of the influence of stenosis severity on the flow in the ascending aorta. Med Eng Phys. 2014;36:1147–55. doi:10.1016/j.medengphy.2014.06.018.

5. Glor FP, Westenberg JJM, Vierendeels J, Danilouchkine M, Verdonck P. Validation of the coupling of magnetic resonance imaging velocity measurements with computational fluid dynamics in a U bend. Artif Organs. 2002;26:622–35. doi:10.1046/j.1525-1594.2002.07085.x.

6. Faghih MM, Sharp MK. Modeling and prediction of flow-induced hemolysis: a review. Biomech Model Mechanobiol. 2019;18:845–81. doi:10.1007/s10237-019-01137-1.

7. Miyazaki S, Itatani K, Furusawa T, Nishino T, Sugiyama M, Takehara Y, Yasukochi S. Validation of numerical simulation methods in aortic arch using 4D Flow MRI. Heart Vessels. 2017;32:1032–44. doi:10.1007/s00380-017-0979-2.

8. Numata S, Itatani K, Kanda K, Doi K, Yamazaki S, Morimoto K, et al. Blood flow analysis of the aortic arch using computational fluid dynamics. Eur J Cardiothorac Surg. 2016;49:1578–85. doi:10.1093/ejcts/ezv459.

9. Steinman DA, Thomas JB, Ladak HM, Milner JS, Rutt BK, Spence JD. Reconstruction of carotid bifurcation hemodynamics and wall thickness using computational fluid dynamics and MRI. Magn Reson Med. 2002;47:149–59. doi:10.1002/mrm.10025.

10. Soudah E, Casacuberta J., Gamez-Montero Pj, Pérez Js, Rodríguez-Cancio M, Raush G, et al. Estimation of wall shear stress using 4D Flow cardiovascular MRI and computational fluid dynamics. J. Mech. Med. Biol. 2017;17:1750046. doi:10.1142/S0219519417500464.

11. Ferdian E, Suinesiaputra A, Dubowitz DJ, Zhao D, Wang A, Cowan B, Young AA. 4DFlowNet: Super-Resolution 4D Flow MRI Using Deep Learning and Computational Fluid Dynamics. Front. Phys. 2020. doi:10.3389/fphy.2020.00138.

12. Youssefi P, Gomez A, Arthurs C, Sharma R, Jahangiri M, Alberto Figueroa C. Impact of Patient-Specific Inflow Velocity Profile on Hemodynamics of the Thoracic Aorta. J Biomech Eng 2018. doi:10.1115/1.4037857].

13. Pirola S, Jarral OA, O’Regan DP, Asimakopoulos G, Anderson JR, Pepper JR, et al. Computational study of aortic hemodynamics for patients with an abnormal aortic valve: The importance of secondary flow at the ascending aorta inlet. APL Bioeng. 2018;2:26101. doi:10.1063/1.5011960.

14. Rodgers JL, Jones J, Bolleddu SI, Vanthenapalli S, Rodgers LE, Shah K, et al. Cardiovascular Risks Associated with Gender and Aging. J Cardiovasc Dev Dis 2019. doi:10.3390/jcdd6020019.

15. Benjamin EJ, Muntner P, Alonso A, Bittencourt MS, Callaway CW, Carson AP, et al. Heart Disease and Stroke Statistics-2019 Update: A Report From the American Heart Association. Circulation. 2019;139:e56–e528. doi:10.1161/CIR.0000000000000659.

16. Yushkevich PA, Piven J, Hazlett HC, Smith RG, Ho S, Gee JC, Gerig G. User-guided 3D active contour segmentation of anatomical structures: significantly improved efficiency and reliability. Neuroimage. 2006;31:1116–28. doi:10.1016/j.neuroimage.2006.01.015.

17. Abraham F, Behr M, Heinkenschloss M. Shape optimization in steady blood flow: a numerical study of non-Newtonian effects. Comput Methods Biomech Biomed Engin. 2005;8:127–37. doi:10.1080/10255840500180799.

18. Manchester EL, Pirola S, Salmasi MY, O’Regan DP, Athanasiou T, Xu XY. Analysis of Turbulence Effects in a Patient-Specific Aorta with Aortic Valve Stenosis. Cardiovasc Eng Technol. 2021;12:438–53. doi:10.1007/s13239-021-00536-9.

19. Benim AC, Nahavandi A, Assmann A, Schubert D, Feindt P, Suh SH. Simulation of blood flow in human aorta with emphasis on outlet boundary conditions. Applied Mathematical Modelling. 2011;35:3175–88. doi:10.1016/j.apm.2010.12.022.

20. Morbiducci U, Ponzini R, Gallo D, Bignardi C, Rizzo G. Inflow boundary conditions for image-based computational hemodynamics: impact of idealized versus measured velocity profiles in the human aorta. J Biomech. 2013;46:102–9. doi:10.1016/j.jbiomech.2012.10.012.

21. Papathanasopoulou P, Zhao S, Köhler U, Robertson MB, Long Q, Hoskins P, et al. MRI measurement of time-resolved wall shear stress vectors in a carotid bifurcation model, and comparison with CFD predictions. J. Magn. Reson. Imaging. 2003;17:153–62. doi:10.1002/jmri.10243.

22. Morbiducci U, Ponzini R, Rizzo G, Cadioli M, Esposito A, Cobelli F de, et al. In vivo quantification of helical blood flow in human aorta by time-resolved three-dimensional cine phase contrast magnetic resonance imaging. Ann Biomed Eng. 2009;37:516–31. doi:10.1007/s10439-008-9609-6.

23. Garcia J, van der Palen RLF, Bollache E, Jarvis K, Rose MJ, Barker AJ, et al. Distribution of blood flow velocity in the normal aorta: Effect of age and gender. J. Magn. Reson. Imaging. 2018;47:487–98. doi:10.1002/jmri.25773.

24. Itatani K, Sekine T, Yamagishi M, Maeda Y, Higashitani N, Miyazaki S, et al. Hemodynamic Parameters for Cardiovascular System in 4D Flow MRI: Mathematical Definition and Clinical Applications. Magn Reson Med Sci. 2022;21:380–99. doi:10.2463/mrms.rev.2021-0097.

25. von Knobelsdorff-Brenkenhoff F, Karunaharamoorthy A, Trauzeddel RF, Barker AJ, Blaszczyk E, Markl M, Schulz-Menger J. Aortic flow and wall shear stress in aortic stenosis is associated with left ventricular remodeling. J Cardiovasc Magn Reson 2016. doi:10.1186/1532-429X-18-S1-Q57.

26. Kilner PJ, Yang GZ, Mohiaddin RH, Firmin DN, Longmore DB. Helical and retrograde secondary flow patterns in the aortic arch studied by three-directional magnetic resonance velocity mapping. Circulation. 1993;88:2235–47. doi:10.1161/01.cir.88.5.2235.

27. Markl M, Draney MT, Hope MD, Levin JM, Chan FP, Alley MT, et al. Time-resolved 3-dimensional velocity mapping in the thoracic aorta: visualization of 3-directional blood flow patterns in healthy volunteers and patients. J Comput Assist Tomogr. 2004;28:459–68. doi:10.1097/00004728-200407000-00005.

28. Torii R, Wood NB, Hadjiloizou N, Dowsey AW, Wright AR, Hughes AD, et al. Fluid–structure interaction analysis of a patient-specific right coronary artery with physiological velocity and pressure waveforms. Commun. Numer. Meth. Engng. 2009;25:565–80. doi:10.1002/cnm.1231.

